# The fat-body secreted neuropeptide CCHa2 signals insulin-producing cells in the brain to promote sleep

**DOI:** 10.1101/2025.11.25.690337

**Authors:** Samaneh Biglari, Shangyu Gong, Miranda C. Wang, Ciara A. Payne, Matthew R. Kauffman, Justin R. DiAngelo, Alex Keene

## Abstract

Sleep is a fundamental behavior regulated by diverse environmental and physiological cues. While the central mechanisms underlying sleep regulation have been investigated in detail, far less is known about how nutrient signals from the periphery are communicated to the brain to modulate sleep. Here, we perform a targeted RNAi screen of genes enriched in the fat body and regulated by feeding state to identify genes that function in the fat body to regulate sleep in *Drosophila*. This analysis found that body-specific knockdown of CCHa2 significantly reduced sleep duration and sleep depth in fed flies, phenocopying sleep in starved flies. *CCHa2*-deficient flies exhibited reduced glycogen stores and diminished feeding drive. Analysis of single-cell transcriptomic atlases confirms that the CCHa2 receptor (CCHa2-R) is selectively expressed in Insulin Producing Cells (IPCs) of the fly brain. We find that knockdown of *CCHa2*-*R* in IPCs recapitulates the sleep loss phenotype of *CCHa2* mutants, supporting a role in adipose-brain signaling. Further screening of genes identified in single cell atlases as being enriched in IPCs led to the identification of numerous genes that function in IPCs to regulate sleep including modulators of *wnt* and insulin signaling. Together, these findings identify a fat body-IPC axis that is a critical modulator of sleep duration and intensity.

## Introduction

Sleep is critical for maintaining essential biological processes including brain function, and immune response, physiology, and memory [1]. Despite its essential role in diverse processes, animals modulate sleep duration and timing in response to diverse environmental factors including light, diet, temperature, stress, and social interactions [2]. Growing evidence suggests distinct cellular processes regulate sleep across different contexts [3]. Identifying the mechanistic basis of sleep regulation under these varying conditions remains a major challenge and is essential for understanding how environmental factors interact to influence sleep states and contribute to sleep-related pathologies.

Flies, rodents, and humans all suppress sleep or experience diminished sleep quality when deprived of food [4–8]. Metabolic regulation of sleep involves nutrient-sensing neurons in the brain that detect changes in circulating nutrients. In mammals, glucose-sensitive neurons have been identified in the hypothalamus that are critical for sleep regulation, and food-regulatory neurons are known to regulate sleep [9,10]. In the fruit fly, *Drosophila*, a single pair of Lateral Horn Leucokinin neurons are activated by circulating nutrient levels and required for metabolic regulation of sleep [11]. Similarly in mammals, shared populations of neurons modulate both sleep and feeding, supporting the notion that there are strong functional interactions between these behaviors [10,12].

Sleep is also potently regulated by peripheral cues from numerous different organs that signal both feeding state and energy stores [13–16]. Adipose tissue senses nutrient levels within the body and modulates hunger-induced behaviors through metabolic control of energy storage, as well as through secreted factors that regulate neural control of behavior [17–19]. Communication between adipose tissue and the brain has received considerable attention because sleep loss is associated with increased feeding, and growing evidence suggests adipose tissue signals nutrient availability to the brain to modulate sleep [20,21]. Identifying the mechanisms of communication between the brain and peripheral tissues is essential for understanding how physiological state modulates sleep.

*Drosophila* is a leading model for studying genetic regulation of sleep and metabolism [22,23]. Mechanisms underlying each of these processes are highly conserved from flies to mammals [22,24–26]. The *Drosophila* fat body is central to the control of energy homeostasis and is the primary site of glycogen and triglyceride storage, thus serving functions analogous to mammalian adipose tissue [27]. Several lines of evidence indicate that adipose tissue and the fat body impact sleep [15,28,29]. First, the *Drosophila* fat body secretes multiple leptin analogs, including the cytokine-like factors *unpaired 1* (*Upd1)* and *unpaired 2* (*upd2)* that act on the brain to modulate feeding [30,31]. Given the role of mammalian leptin in sleep regulation, it is possible that *upd1/2* or other secreted factors impact sleep. Second, mutations of numerous genes in the fat body including *lipid storage droplet 2* (*lsd2*), *Brummer*, and *ade2* alter sleep regulation [6,15]. Therefore, targeted screening of the fat body has potential to uncover novel sleep regulatory mechanisms.

Here, we perform a genetic screen by selectively knocking down genes in the fat body and measuring the effect on sleep. We specifically targeted genes with transcripts that were differentially expressed in accordance with feeding state, or genes whose products are predicted to be secreted from the fat body. This screen identified numerous genes that function in the fat body to regulate sleep, including the fat-body secreted neuropeptide CCHa2. Together, these findings identify novel adipose-derived factors that regulate sleep and contribute to adipose-brain communication.

## Results

To identify genes that function in the fat body to regulate sleep, we first selected candidates based on prior evidence of differential expression during sleep and feeding states, enrichment in the fat body, and predicted secretion (Table 1) [32,33]. In total, 81 genes with available RNAi lines were identified and screened using the fat body-specific driver CG-GAL4 to selectively knock down genes in the fat body (Fig 1A) [34,35]. Flies harboring both an RNAi transgene targeting a candidate gene and CG-GAL4 were compared to control flies harboring CG-GAL4 crossed to the genetic background strain from each RNAi library, as well as CG-GAL4 crossed to Luciferase-RNAi [36,37]. In total, this screen identified 19 RNAi lines with sleep that was significantly reduced compared to controls, and sleep was significantly increased in 7 RNAi lines (Fig 1B). This screen identified numerous genes known to regulate sleep including sleep-promoting roles for the cytokine unpaired 2 (*upd2*) and the lipid-storage droplet-associated lipase *brummer* [32,38]. Several genes that had not been previously implicated in sleep regulation were also identified including the stress-inducible *apolipoprotein neuropeptide-like precursor 2* (NPLP2) and the Jak/Stat target *turnadot* C (*TotC*)[39,40].

**FIGURE 1.**
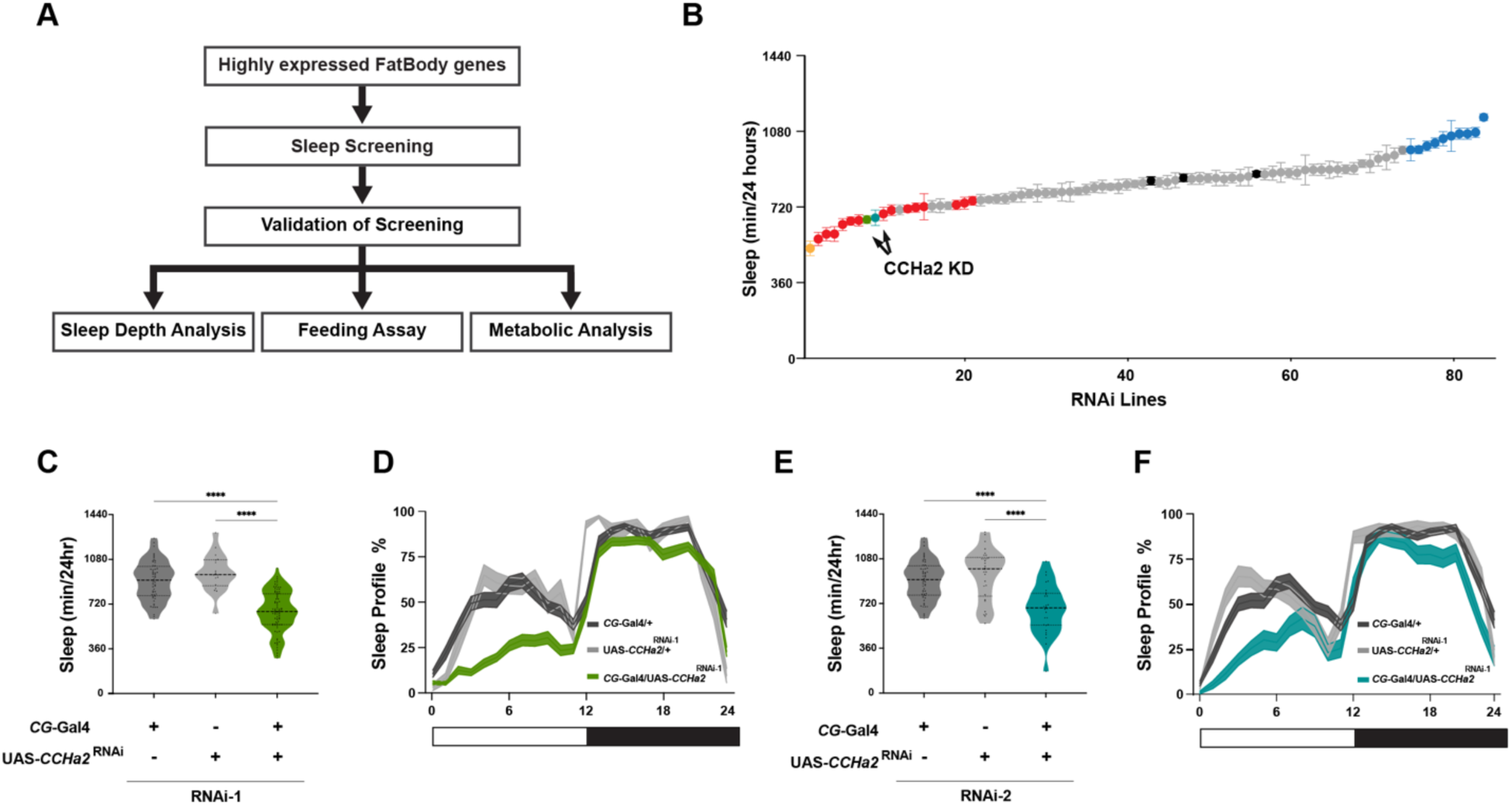
*CCHa2* knockdown leads to sleep loss. (A) Schematic of the screening strategy used to identify fat body-expressed genes regulating sleep. (B) Total sleep (minutes over 24 hours) across viable RNAi lines (81 lines, n > 32 per line). Error bars indicate the SEM. Black dots indicate control sleep values; green and teal dots highlight RNAi lines targeting *CCHa2* that fall outside the control range; orange dot shows the lipid-storage droplet-associated lipase *brummer*, red and blue dots show sleep responses of RNAi lines that fall outside the control range (C) Total sleep is significantly reduced in *CCHa2* RNAi line compared to controls (*CCHa2* RNAi1, green). (D) Knockdown of *CCHa2* significantly reduces daytime sleep. (One-way ANOVA: F _2, 156_ =54.59, \*\**p* < 0.0001). (E-F) A second, independently derived RNAi line validated the same sleep loss phenotype (*CCHa2* RNAi2, teal). (F _2, 111_ = 21.00, \*\**p* < 0.0001). Lines and error bars represent mean ± SEM; Horizontal lines represent quartiles. Asterisks indicate statistically significant differences between groups.

We chose to focus further analysis on the gene *CCHa2* because it encodes an appetite-regulating neuropeptide that may be secreted from the fat body to promote sleep [41,42]. Previous studies have examined the role of CCHa2 secretion from the gut, but its role in sleep regulation has not been investigated [41,42]. To confirm findings from the screen, *CCHa2* knockdown flies were retested with additional heterozygous controls. Flies with RNAi targeted to *CCHa2* (*CG-*GAL4*>CCHa2-*RNAi) slept significantly less than flies harboring *CG-*GAL4 or *CCHa2*-RNAi alone (Fig 1C, D). To validate that the findings were not caused by off-target effects, we assayed a second, independently derived RNAi line. Flies with a second RNAi line targeting *CCHa2* <u>to the fat body</u> (*CG-*GAL4>*CCHa2-*RNAi^2^) phenocopy the <u>sleep loss phenotype</u> of the <u>first RNAi line</u> (Fig 1E, F). For both RNAi lines, sleep was significantly reduced in *CG-*GAL4*>CCHa2-*RNAi during the daytime (Fig S1A, B). Therefore, these results suggest that fat-body derived CCHa2 promotes sleep.

The *CG-*GAL4 driver is expressed in the fat body but may also be expressed in other tissue types during adulthood and across development [43,44]. To validate that fat body-specific knockdown of *CCHa2* inhibits sleep, we repeated experiments using the fat body driver *r4-*GAL4 [45]. Flies harboring *r4-*GAL4*>CCHa2-*RNAi slept less than heterozygous controls during both the day and night, phenocopying the sleep phenotype observed with *CG-*GAL4 knockdown (Fig. S1C). Therefore, multiple independent GAL4 drivers and RNAi lines confirm a sleep-promoting role for fat body CCHa2.

Fly sleep can be divided into light and deep sleep, each of which has distinct behavioral and physiological characteristics [46]. Hidden Markov Models have been applied to quantify the probability of maintaining a waking state p(Wake) or a sleeping state, termed p(Doze), which is proposed to be a measure of sleep depth (Fig 2A) [47]. We found that wake propensity P(Wake) was elevated in both *CG-*GAL4*>CCHa2-*RNAi lines tested, compared to each set of controls (Fig 2B). Conversely, sleep propensity p(Doze) was reduced in both RNAi lines (Fig 2C). There were no significant differences in P(Wake) or P(Doze) during nighttime sleep (Fig S2A,B).

**FIGURE 2.**
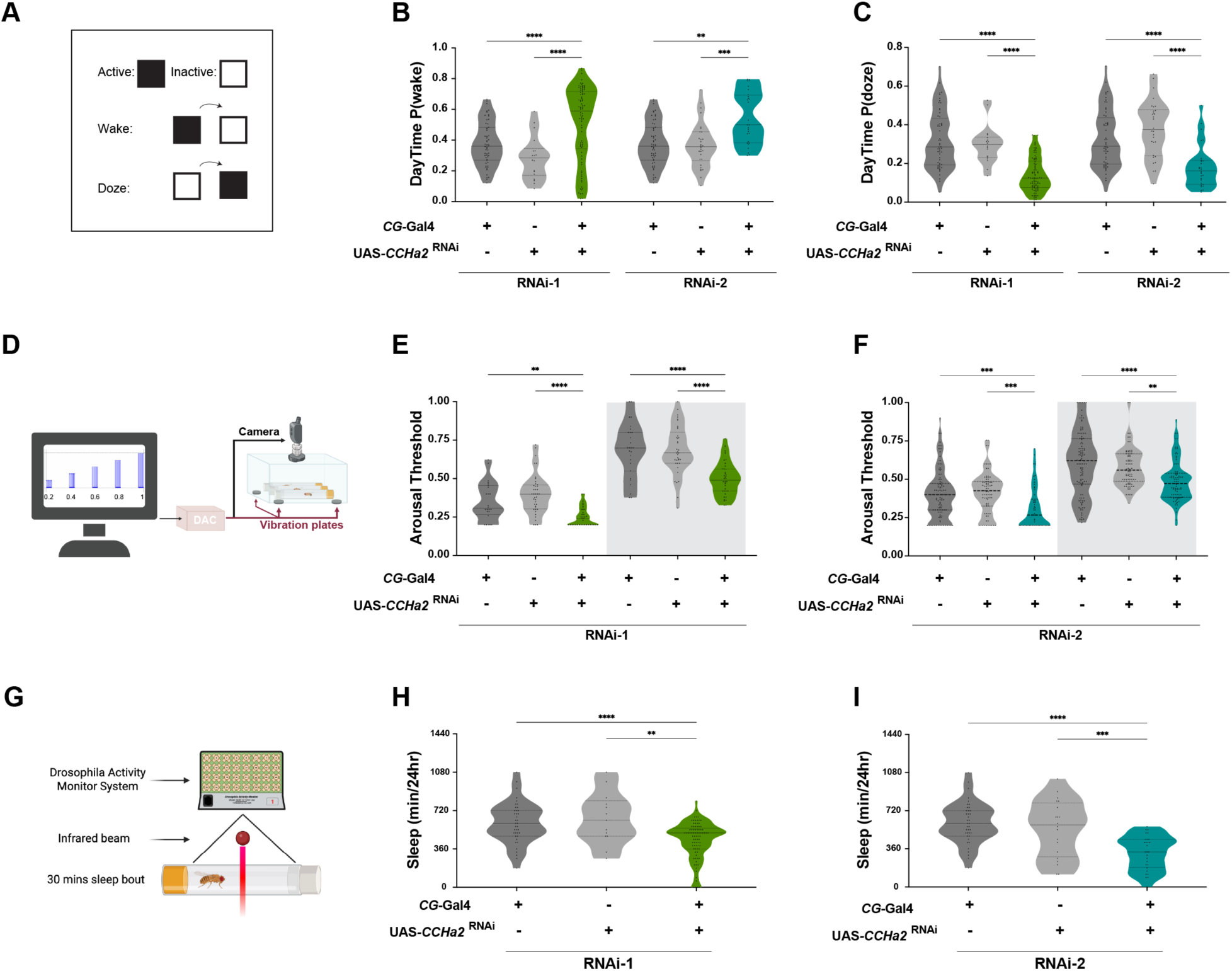
*CCHa2* knockdown decreases sleep depth. (A) Activity pattern of Hidden Markov Model (HMM). Dark squares indicate inactivation, and white squares show activation. (B) HMM analysis reveals that the probability of waking (P(Wake)) is significantly increased in both *CCHa2* RNAi lines compared to controls. Statistical analysis: Kruskal–Wallis test followed by Dunn’s multiple comparisons test (α = 0.05). RNAi^1^: \*\**p* < 0.0001 n = 86 (*CG*/*CCHa2* RNAi), 58 (*CG*/+), and 16 (+/*CCHa2*RNAi). RNAi^2^: \*\**p* < 0.0001 n = 28 (*CG*/*CCHa2*RNAi), 58 (*CG*/+), and 30 (+/*CCHa2*RNAi). (C) The probability of dozing (P(Doze)) is significantly reduced in both *CCHa2* RNAi lines. RNAi: *p* < 0.0001, n = same as above. (D) The *Drosophila* Arousal Threshold (DART) assay measures sleep depth by recording every movement of flies while a digital analog converter (DAC) controls mechanical stimuli. Mechanical stimuli are applied to three platforms, each containing twenty flies, controlled by two motors. Stimuli of progressively increasing intensity are delivered to evaluate arousal thresholds, which are recorded by the computer system. Measurements of arousal threshold are conducted hourly, beginning at Zeitgeber time (ZT) 0. (E-F). Arousal threshold is significantly reduced in RNAi^1^ during both daytime (F_2,98_ = 18.37, **P<0.0001) and nighttime (F_2,99_ = 19.85, P<0.0001) and in RNAi^2^ during both phases as well (Day: F_2,233_ = 10.96, P<0.0001; Night: F_2,273_ = 12.92, P<0.0001). (G) To measure deep sleep, we analyzed for total sleep bouts lasting 25 minutes or longer, (H-I) Total sleep was reanalyzed using a 30-minute minimum bout length as an alternative threshold to confirm the reduction in deep sleep, validating the phenotype observed with the conventional 5-minute threshold. (One-way ANOVA: F _2, 134_ =13.74, *p* < 0.0001, F _2, 84_ = 16.01, *p* < 0.0001). Lines and error bars represent mean ± SEM; Horizontal lines represent quartiles. Asterisks indicate significant differences.

To further examine sleep intensity, we tested flies in the *Drosophila* ARousal Threshold (DART) assay, which has been widely used to quantify sleep duration and intensity [46,48]. During deep sleep, the arousal threshold is elevated in flies, requiring greater mechanical stimulus to induce activity (Fig 2D). For both RNAi lines tested, arousal threshold was significantly reduced during both the daytime and nighttime compared to controls (Fig 2E,F). We also analyzed sleep for total sleep bouts lasting 25 minutes or longer, a measure that is predicted to represent deep sleep (Fig 2G) [49]. For both RNAi lines, fat body-specific knockdown of *CCHa2* resulted in significantly less sleep than control flies harboring *CG-*Gal4 or *CCHa2-*RNAi alone (Fig 2H,I). Together, these findings confirm that sleep intensity is reduced in flies deficient for fat body *CCHa2*.

To examine whether loss of *CCHa2* in the fat body results in metabolic dysregulation, we measured energy stores in flies under fed and starved conditions (Fig 3A). Under fed conditions glycogen stores were significantly reduced in both the *CG-*GAL4>*CCHa2*-RNAi (Fig 3B). No differences were detected in free glucose levels for either RNAi lines. Triglyceride level was increased with only one of the RNAi lines (Fig S3A,B). Together, these findings reveal deficits in energy store accumulation in *CCHA2*-defficient flies.

**FIGURE 3.**
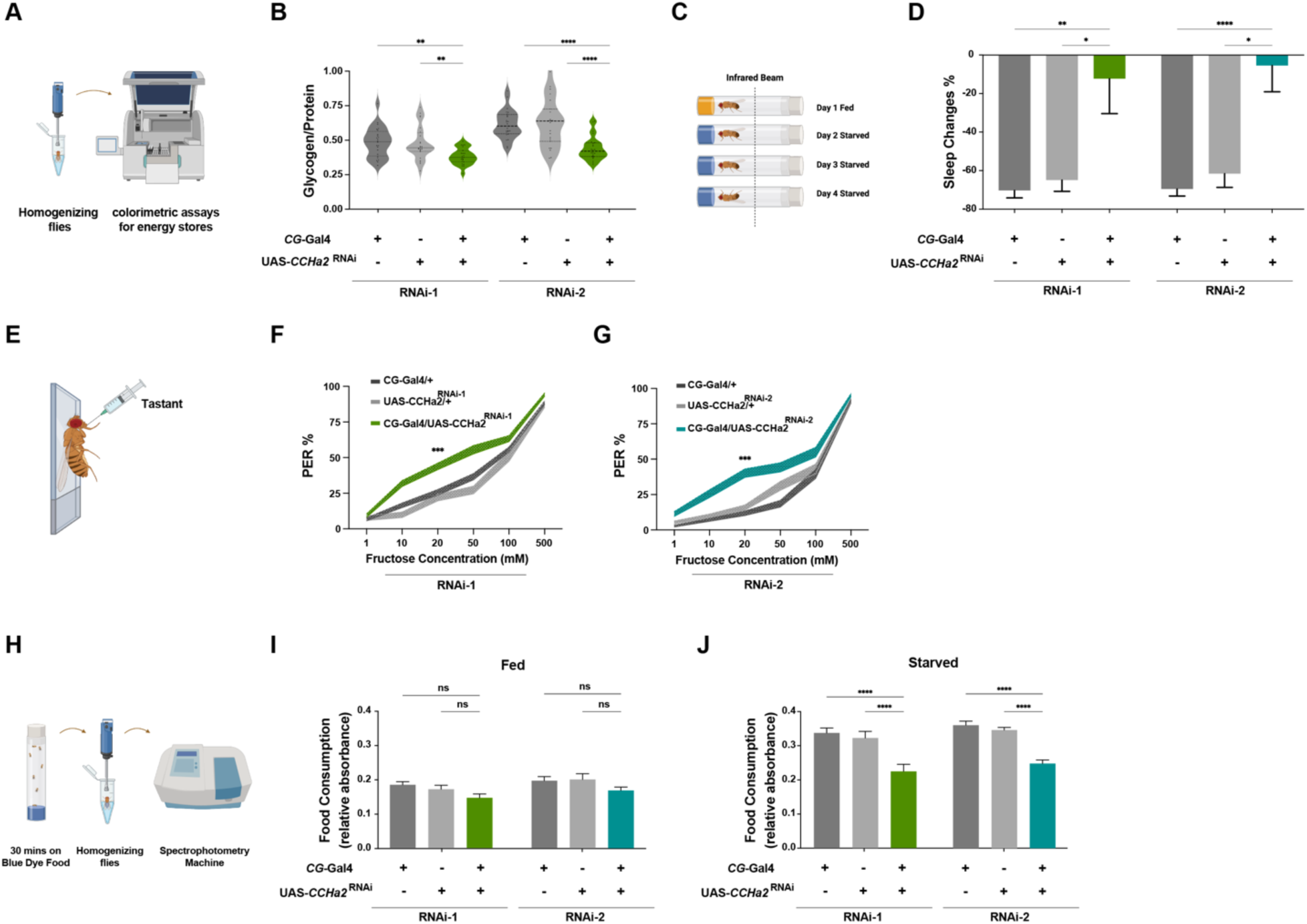
*CCHa2* knockdown affects metabolic and behavioral responses to starvation. (A) Triglyceride, glycogen, free glucose, and protein levels are measured by homogenizing two female fly bodies (3–5 days old) in a standard buffer (50 mM Tris-HCl (pH 7.4), 140 mM NaCl, 0.1% Triton-X, and 1X protease inhibitor cocktail), and colorimetric assays are used to check the energy stores. (B) Both *CCHa2* RNAi line flies show reduced Glycogen levels under fed conditions, with no significant difference under starvation (N=24; *p* > 0.0010 and *p* > 0.1060, respectively). (C) Flies are monitored for sleep over a 24h period on standard food, then transferred to agar and maintained until death to evaluate both starvation-induced sleep suppression and starvation resistance. (D) Starvation-induced sleep suppression, measured as percentage change relative to the fed condition, is significantly decreased in *CCHa2* knockdown flies compared to controls (*p* < 0.0001 for all comparisons). (E) Schematic of the proboscis extension reflex (PER) assay. Sugar is applied to the labellum of a tethered female fly. (F-G) After 24 hours of starvation, PER is significantly increased in both *CCHa2* RNAi lines at 20 mM sucrose (n > 30; RNAi^1^: *p* = 0.005936, RNAi^2^ p = 0.006948). (H) Starved or fed flies were transferred to blue dye food for 30 minutes. After homogenizing them, spectrophotometry was used to quantify the amount of consumed food. (I-J) Spectrophotometric quantification determines significantly increased food consumption in starved control flies. Two-way ANOVA revealed a significant interaction genotype in nutritional state interaction (RNAi^1^: F_2, 98_= 4.261, *p* = 0.0168, RNAi^2^: F_2, 139_= 8.348, *p* < 0.0004), as well as main effects of genotype (RNAi^1^: F_1, 98_ = 104.7, *p* < 0.0001, RNAi^2^: F_1, 139_ = 185.1, *p* < 0.0001) and nutritional condition (RNAi^1^: F_2, 98_ = 14.99, *p* < 0.0001, RNAi^2^: F_2, 139_ = 26.55, *p* < 0.0001). Lines and error bars represent mean ± SEM.

Flies suppress their sleep when starved, presumably to increase time spent foraging [5]. Given the role of the fat body in sleep regulation, we sought to determine whether CCHa2 is involved in modulating sleep in accordance with nutrient availability [15,50]. We measured starvation-induced sleep suppression, comparing sleep for 24 hours on standard fly food to 24 hours under starvation conditions where flies were transferred to tubes containing agar (Fig 3C). In control flies, sleep was greater on food than on agar (Fig 3D). Conversely, *CG-*GAL4*>CCHa2-*RNAi knockdown flies did not suppress sleep during starvation (Fig 3D; Fig S3C-F). Furthermore, under starved conditions, sleep in *CCHa2*-knockdown flies did not differ significantly from controls (Fig S3C-F). Together, these findings reveal a loss in diet-dependent modulation of sleep in *CCHa2*-knockdown flies.

Feeding drive is increased during starvation [51,52]. When presented with an appetitive tastant, flies exhibit a Proboscis Extension Reflex (PER), providing a quantifiable measure of feeding propensity (Fig 3E) [53]. To determine if reflexive feeding response is increased in *CCHa2-*RNAi flies, we measured PER in response to presentation of appetitive fructose at concentrations ranging from 1-500 mM. At a moderate concentration of 20 mM, PER was increased in both *CG-*GAL4*>CCHa2-*RNAi lines compared to heterozygous controls (Fig 3F,G). At higher concentrations there was no difference between *CG-*GAL4*>CCHa2-*RNAi and controls, indicating that the result is not due to developmental impairments in PER. These findings reveal enhanced PER in response to tastants.

To determine whether fat body-specific knockdown of *CCHa2* influences the amount of food consumed, we compared *CG-*GAL4*>CCHa2-*RNAi flies with heterozygous controls using the blue dye assay. Briefly, fed or 24hr starved flies were housed on food containing blue-dye for 30 minutes [54,55]. At the completion of this assay flies were flash frozen and spectrophotometry was used to quantify the amount of food consumed (Fig 3H). There were no differences between *CG-*GAL4*>CCHa2-*RNAi flies and controls during the fed state (Fig 3I). However, in starved flies, food consumption was reduced in *CCHa2-*RNAi knockdown flies compared to controls (Fig 3J). Together, these findings suggest fat body-specific CCHa2 promotes hunger-induced increases in food intake.

The *Drosophila* genome encodes for a single CCHa2 receptor (CCHa2R) that regulates the transcription of *Drosophila insulin-like peptides* (*ilps*) [56]. To examine where *CCHa2-R* is expressed across the body, we analyzed *CCHa2* and *CCHa2-R* expression merging multiple *Drosophila* single-cell atlases [57,58]. Analysis confirmed that *CCHa2* is highly expressed in the fat body, while *CCHa2-R* is predominantly restricted to the Insulin Producing Cells (IPCs) in the brain (Fig 4A,B). The IPCs are critical regulators of sleep and are targets of both fat-body secreted *upd2* and the neuropeptide Leucokinin that is critical for starvation-induced sleep suppression [59–61]. To test if *CCH<u>a</u>2-R* functions in the IPCs to regulate sleep, we targeted expression to all neurons <u>using a pan-neuronal GAL4</u> (*Nsyb-GAL4*), or selectively in the IPCs under control of *ilp2-*GAL4. Pan-neuronal knockdown of *CCHa2-R* with *Nsyb-*GAL4, or knockdown with *ilp2-*GAL4 significantly reduced sleep phenocopying *CCHa2* knockdown (Fig 4C). Sleep was significantly reduced in *ilp2-*GAL4*>CCHa2-R-RNAi* during both daytime and nighttime (Fig 4D,E). No effect was observed in flies with *CCHa2-R* knockdown in dopamine neurons (*TH-*GAL4) that regulates sleep and feeding through non-overlapping populations of neurons (Fig S4A).

**FIGURE 4.**
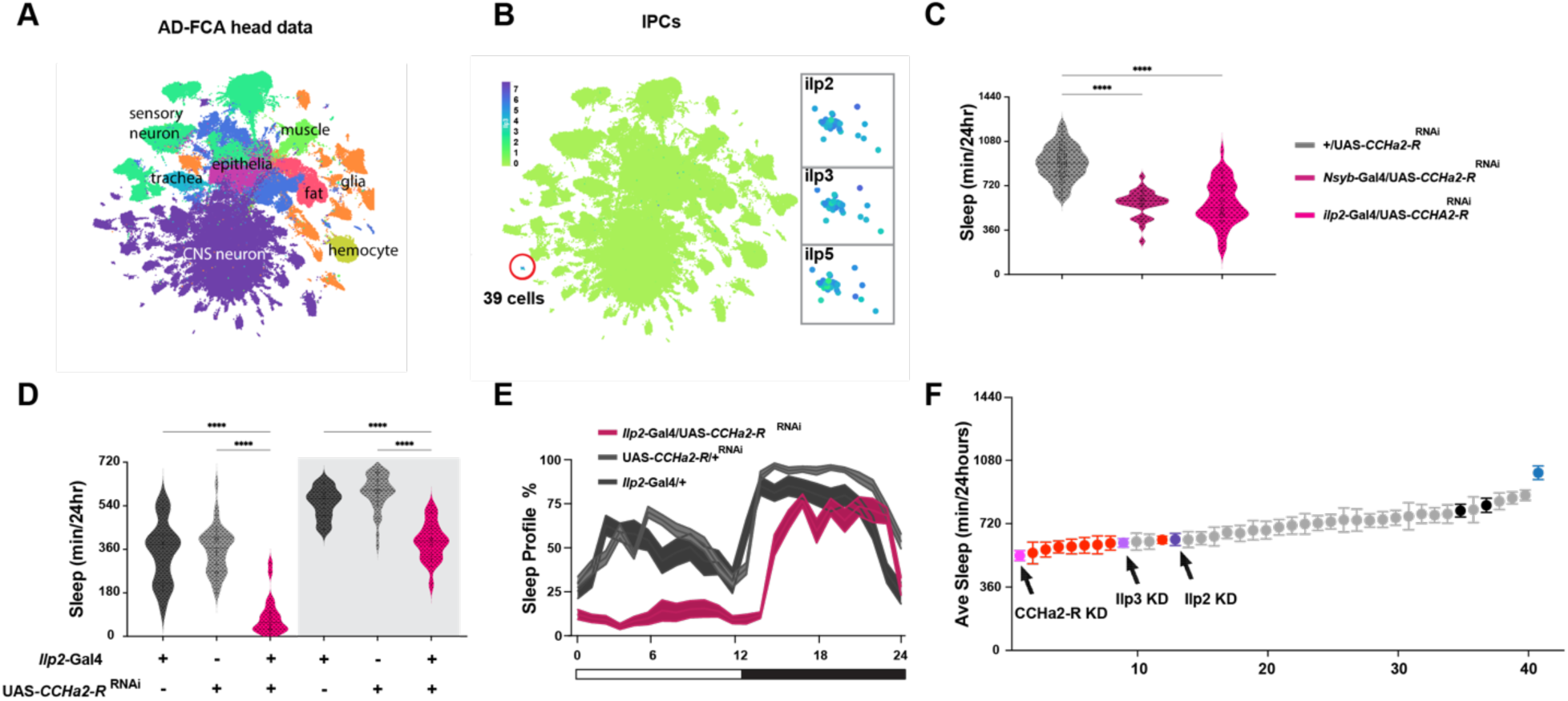
*CCHa2-R* knockdown in neurons reduces sleep and alters sleep architecture. (A) Alzheimer’s Disease Fly Cell Atlas (AD-FCA) head data is used to determine which cell-types CCHa2-R is mainly expressed in. (B) Analysis revealed that CCHa2-R is predominantly restricted to the Insulin Producing Cells in the brain. (C) Pan-neuronal (*Nsyb-*GAL4) and *ilp2*-specific knockdown of *CCHa2-R* significantly reduces total sleep. (*p* <0.0001). (D) Knockdown of *CCHa2-R* in IPC significantly decreases both daytime (F_2, 75_= 60.63, *p* <0.0001) and nighttime sleep (F_2, 75_= 55.57, *p* <0.0001). (E) Sleep profile shows a dramatic reduction in both phases. Lines and error bars represent the mean ± SEM. (F) RNAi-based screening of genes highly expressed in IPC (*ilp2-*GAL4 driver) reveals that knockdown of *Ilp3* (light purple) and *Ilp2* (dark purple) significantly reduced total sleep (42 lines, n > 16 per line). Black dots indicate controls. Pink dot show CCHa2R. Red and blue dots show sleep responses of RNAi lines that fall outside the control range. Error bars indicate the SEM.

To confirm these findings, we applied Hidden Markov Modeling and arousal threshold measurements. Wake propensity was increased, and sleep propensity was decreased in flies with loss of *CCHa2-R* in the IPCs, phenocopying fat body specific knockdown of *CCHa2* (Fig S4B,C). Similarly, arousal threshold of *ilp2-*GAL4*>CCHa2R-*RNAi was significantly reduced during the daytime and nighttime compared to control flies (Fig S4D). While mean sleep bout length in *CCHa2-R* knockdown flies was not significantly altered compared to controls, bout number was significantly reduced, indicating a net decrease in total sleep (Fig S4E,F). Together, these findings reveal that CCHa2R functions in the IPCs to promote sleep duration and increase sleep depth.

To identify genes that function within the IPCs to regulate sleep, we mined snRNA-seq data sets for transcripts that are upregulated in CCHa2-R-expressing IPCs [57,58]. From this analysis, we screened 40 genes that were highly expressed in the IPCs (Table 2) [57,58]. We then expressed RNAi targeting each of these genes under-control of *ilp2-*GAL4 [62]. Knockdown of <u>each of</u> these 40 genes revealed 10 genes with reduced sleep including *ilp2* and *ilp3*, led to reduced sleep compared to *ilp2-*GAL4/+ control flies, consistent with a previously reported sleep-promoting role form *ilps* (Fig S4G,H) [59]. We also identified numerous additional genes with shorter sleep than control flies including the inhibitor of *wnt* signaling *naked cuiticle* (*nkd)*, *insulin growth factor* II mRNA *binding protein* (*imp*), the *homeobox transcription factor* (*ey*) and the cell adhesion molecule *beaten path 1b* (*beat-1B)* (Fig 4F). These findings suggest IPCs play a critical sleep promoting role, and that *ilp2/3* output in response to fat body secreted *CCHa2* modulates sleep.

## Discussion

In mammals, feeding state potently regulates sleep, primarily through the modulation of orexigenic neurons in the hypothalamus [63]. The permeability of the blood-brain barrier is modulated by hypothalamic neurons and there is evidence that adipose-secreted molecules, including leptin contribute to sleep regulation [64,65]. The *Drosophila* fat body is the functional analog of the mammalian adipose tissue, and these findings build on prior studies suggesting a sleep-regulating axis between the fat body and modulatory neurons in the brain that promote sleep. Numerous genes have been identified that function within the fat body to regulate sleep including a role for the triglyceride lipase *brummer* (*bmm*) in modulating sleep architecture, and in the purine biosynthesis pathway member *Ade2,* in regulating both energy stores and suppressing sleep [6,15]. Beyond intracellular metabolism, the fat body communicates with the brain via secreted signals [66]. Notably, *unpaired 2* (*upd2*), a leptin-like cytokine, is secreted from the fat body in response to nutrient status and modulates sleep by signaling through the JAK/STAT pathway in the brain[29]. Our findings add to these previous studies and suggest the neuropeptide CCHa2 is secreted from the fat body to modulate sleep. The finding that numerous fat body-secreted genes and signaling molecules modulate sleep reveals dynamic interactions between the fat body and the brain that have largely been overlooked.

The neuropeptides CCHa1 and CCHa2 regulate behavioral and physiological responses to changes in nutrient availability in flies [41,67]. CCHa1 has been identified as a critical regulator of nutrient-dependent changes in arousal threshold, with loss of *CCHa1* resulting in reduced arousal threshold, similar to our findings in *CCHa2* mutant flies [68]. CCHa1 is particularly responsive to the effects of dietary protein on elevated arousal threshold and starvation-dependent enhancement of sensitivity to food odor [67,68]. We find that loss of *CCHa2* in the fat body also impairs dietary modulation of sleep, though it remains to be determined whether this is due to a general loss of circulating nutrients, or defined macronutrients. Beyond its role in sleep regulation, both CCHa1 and CCHa2 are expressed in the gut and fat body. CCHa1 functions in the gut to broadly regulate nutrient sensing behaviors, including sensory neuron excitability and foraging [68]. CCHa2 supports sustained feeding and growth-related behaviors, linking peripheral nutrient sensing to central control of energy balance [56]. Therefore, both CCHa1 and CCHa2 play critical roles in modulating the response to nutrients and defining how each neuropeptide functions within defined cell types and in response to nutritional availability will be critical for understanding the role of brain-periphery communication in sleep and other behaviors.

Our findings suggest complex roles for fat body CCHa2 in the modulation of taste, feeding, and diet-dependent changes in sleep. A previous study found that feeding is reduced in *CCHa2* mutant larvae and adults, and this effect is partially rescued by restoring CCHa2 to the gut, suggesting an orexigenic role of CCHa2 secretion from the gut [41]. Therefore, it is possible that CCHa2 secretion from both the gut and fat body are required for normal feeding. *CCHa2* mutants were also reported to have reduced locomotor activity, in agreement with our findings [15,69]. However, we also found that reflexive feeding in flies is elevated in flies with loss of *CCHa2* in the fat body suggesting that *CCHa2* uniquely impacts distinct behavioral components of foraging. Together, our findings indicate that the effects of loss of *CCHa2* from the fat body are complex, enhancing sensory responsiveness to a tastant, while reducing overall feeding. Identifying the cell-type specific differences in CCHa2 function, including the full complement of target cells in the brain and periphery will require examining state-dependent changes in transcription, release, and broader analysis of receptor localization.

Our findings suggest that CCHa2 release from the fat body signals IPCs in the brain to modulate sleep. The IPCs are central integrators of nutrient state and sleep regulation that are thought to coordinate input from the brain and periphery to modulate release of *ilps* and other neuropeptides that modulate behavior [70,71]. The physiology and transcriptional regulation of these cells are modulated by the fat-body secreted *upd2*, that in turn signals starvation and suppresses sleep. Further, the Lateral Horn Leucokinin (LHLK) neurons, which increase activity during starvation and are required for integration of sleep and metabolic state, innervate the IPCs [11]. Knockdown of the Leucokinin Receptor (*LKR*) within the IPCs phenocopies silencing of LHLK neurons, suggesting the IPCs are responsive to LHLK neurons [11]. This supports the notion that IPCs are a central integrator of sleep and metabolic state receiving input from the brain and periphery. Beyond direct regulation, sleep loss itself can trigger widespread metabolic changes, including changes in energy mobilization and gut function [72]. IPCs not only regulate sleep but also mediate long-term responses to diet and stress, including effects on aging, neuroprotection, and systemic energy balance. Together these findings suggest the IPCs are a central integrator of sleep and metabolic state receiving input from the brain and periphery.

To date the vast majority of large-scale screens for sleep-regulating genes in fruit flies have either used classic genetic mutation, ubiquitous RNAi knockdown, or RNAi knockdown targeted to the brain [73–77]. Single cell atlases provide a cellular resolution of gene expression, allowing for targeted approaches to identify genes that function in defined cell types. The recent development of single-cell atlases across tissue-types, developmental stages, and in disease models, provides a rich source of information for how IPCs, and other neuromodulatory cells change across context<u>s</u> [57,78].

In humans, sleep exerts a strong influence on glucose and insulin metabolism. Glucose, the brain’s primary energy source, is tightly regulated by insulin, and glucose tolerance decreases during sleep, leading to elevated glucose and insulin levels that peak during sleep [79,80]. This decline in tolerance is largely attributed to reduced glucose utilization by the brain and peripheral tissues in early sleep [80]. In *Drosophila*, insulin-like peptides (ILPs) are essential for metabolic homeostasis and play a role in sleep regulation [81]. Loss of ILP2 or ILP3 reduces sleep depth and duration, indicating a sleep-promoting function [82]. Starvation-induced activation of LKN-producing neurons strongly inhibits insulin-producing cells (IPCs), lowering ILP release and reducing sleep, consistent with this role [11,69]. However, AstA-producing neurons (AstANs), which promote sleep and suppress feeding, also inhibit IPCs, rapidly decreasing ILP release [83,84]. This apparent contradiction suggests that neuromodulatory regulation of sleep and ILP signaling can act through parallel, context-dependent mechanisms.

Further, a recently developed single cell atlas examines gene expression under conditions of sleep deprivation and circadian timepoints [85]. Here, we were able to target specific genes that are expressed in the IPCs, a known population of sleep regulating neurons, to efficiently target candidate genes. Combining the rapidly growing single-cell data, with high-throughput screening has the opportunity to identify novel genes with greater efficiency. Together, these findings identify CCHa2 as a key adipose-derived signal that promotes sleep by acting on IPCs in the brain. Our study reveals a fat body–brain axis through which CCHa2 regulates sleep, presumably by modulating IPC function. Future work examining the effects of CCHa2/CCHa2R signaling on the physiology and transcriptional regulation of IPCs has potential to identify novel mechanisms through which periphery-brain communication modulates sleep. Understanding how peripheral tissues such as the fat body communicate nutrient status to the brain provides critical insight into the mechanisms linking metabolic state to behavioral output, with broad relevance to the regulation of sleep and energy balance across species.

## Methods

### Fly Stocks

Flies were reared on standard cornmeal-based food (Bloomington recipe, Genesee Scientific) and maintained in incubators (Powers Scientific; Dros52) at 25 °C on a 12:12 LD cycle, with relative humidity maintained between 55% and 65%. The *w1118* strain was used as the genetic background control. Unless otherwise noted, all experimental fly lines were outcrossed to this background for 6–8 generations. Fly stocks were obtained from the Bloomington Drosophila Stock Center, including *w1118* (5905; Levis et al. 1985), *CG-*GAL4 (7011; [44], and *r4-*GAL4 (33832; Lee and Park 2004). RNAi lines used in the screen were sourced from the TRiP collection [36,86] and are detailed in Table 1.

### Sleep analysis

Sleep was recorded using the Drosophila Activity Monitor System (DAMS), which tracks fly movement based on infrared beam crossings [87]. Sleep was defined as periods of immobility lasting at least 5 minutes and analyzed using the Drosophila Counting Macro [88,89]. All sleep experiments were conducted under a 12:12 LD cycle. Female flies, aged 5–7 days, were lightly anesthetized with CO_2_ and placed in plastic tubes containing standard food. After a 24-hour recovery period, sleep was monitored for 24 hours (ZT0–ZT24).

### Arousal Threshold Assay

Arousal thresholds were assessed using the Drosophila Arousal Tracking (DART) system as described [46]. Individual female flies were housed in plastic tubes (Trikinetics, Waltham, MA) placed on vibration-enabled trays. Flies were stimulated hourly over a 24-hour period (starting at ZT0) using increasing vibration intensities ranging from 0 to 1.2 g in 0.3 g increments (each lasting 200 ms), with 15-second intervals between stimuli. Flies were continuously recorded at 1 frame per second using a USB webcam (Logitech). Vibration delivery, video acquisition, and data analysis were performed using custom MATLAB software (MathWorks, Natick, MA).

### Protein, glucose, glycogen and triglyceride measurements

Measurement of energy storage molecules was performed as previously described [18]. Briefly, two fly bodies from 3–5-day old females are homogenized in buffer containing 50 mM Tris-HCl (pH 7.4), 140 mM NaCl, 0.1% Triton-X, and 1X protease inhibitor cocktail. Total protein content and triglyceride levels were measured using the Pierce BCA protein assay kit (ThermoFisher, Waltham, MA) and Infinity Triglyceride Reagent (ThermoFisher, Waltham, MA), respectively, according to manufacturer’s instructions. Total glucose was measured using the Pointe Scientific Glucose Oxidase Reagent (ThermoFisher, Waltham, MA) after incubating samples with equal parts 8 mg/mL amyloglucosidase (Sigma-Aldrich, St. Louis, MO) in 0.2M citrate, pH 5.0, for two hours at 37°C. Free glucose was measured using the Pointe Scientific Glucose Oxidase Reagent in undigested samples. Glycogen was calculated by subtracting the free glucose from the total glucose. Glycogen, free glucose and triglyceride levels were normalized by dividing by total protein content.

### Proboscis extension response (PER)

Female flies were starved for 24 hours before PER testing, following previously described protocols [90]. Flies were anesthetized using CO_2_ and mounted on microscope slides (#12-550-15, Fisher Scientific) with only the head and proboscis exposed. After a 60-minute acclimation period in a humidified chamber, flies were offered water until satiated. Flies failing to stop responding within 5 minutes were excluded. A Kimwipe wick (#06–666, Fisher Scientific) saturated with tastant solution (prepared in water) was applied to the tip of the proboscis for 1–2 seconds. Full proboscis extensions were scored as positive responses. Each tastant was presented three times at 1-minute intervals, and PER was calculated as the percentage of positive responses per fly. Experiments were repeated three times per week, and both genotypes and tastant presentation order were randomized.

### Blue-dye feeding assay

Short-term food consumption was measured as previously described [91]. Flies were either starved for 24 or 48 hours on moistened Kimwipes or maintained on standard food. At ZT0, flies were transferred to vials containing 1% agar, 5% sucrose, and 2.5% FD&C Blue Dye No. 1. After 30 minutes of feeding, flies were flash-frozen on dry ice and homogenized individually in 400 μL PBS (pH 7.4; Ambion). Absorbance at 655 nm was measured using a 96-well plate reader (Millipore iMark, Billerica, MA). Background absorbance was corrected by subtracting the mean absorbance of non-dye-fed control flies.

### Statistical Analysis

All data are presented as mean ± SEM. Unless otherwise specified, group comparisons were performed using one-way ANOVA followed by Tukey’s post hoc test. For comparisons between two groups, unpaired two-tailed t-tests were used. Arousal threshold data were analyzed using the non-parametric Kruskal–Wallis test with Dunn’s multiple comparisons post hoc correction. Statistical significance was set at *p* < 0.05. All analyses were conducted using GraphPad InStat (version 6.0).

## Acknowledgements

The authors are grateful for technical support and guidance from Hongjie Li (Baylor). This work was supported by NIH Grant R01 NS131628.

---

**FIGURE S1.**
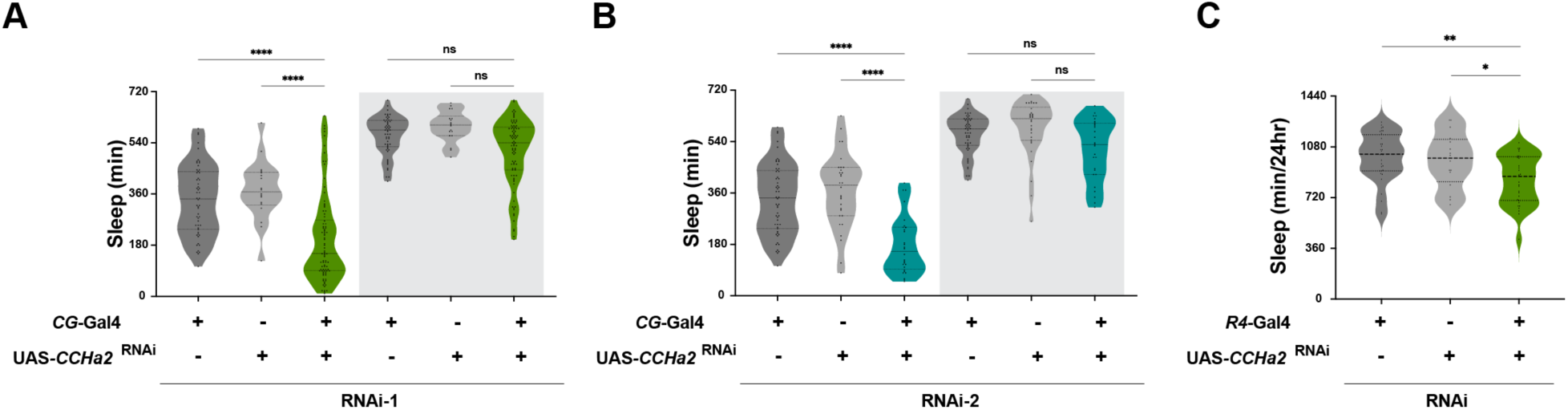
Fat-body drive *CCHa2* promotes sleep. (A-B) For both RNAi lines, sleep is significantly reduced in CCHa2-RNAi KD during the daytime (RNAi^1^: F_5,314_= 6.086, *p* <0.0001, RNAi^2^: F_5,221_= 4.527, *p* <0.0001). (C)Targeted knockdown of *CCHa2* using the fat body-specific driver *r4*-GAL4 (Lee and Park, 2004) significantly reduces total sleep compared to controls, phenocopying results with *CG-*GAL4. One-way ANOVA: F_2, 76_= 6.986, *p* = 0.0016. Lines and error bars represent mean ± SEM.

**FIGURE S2.**
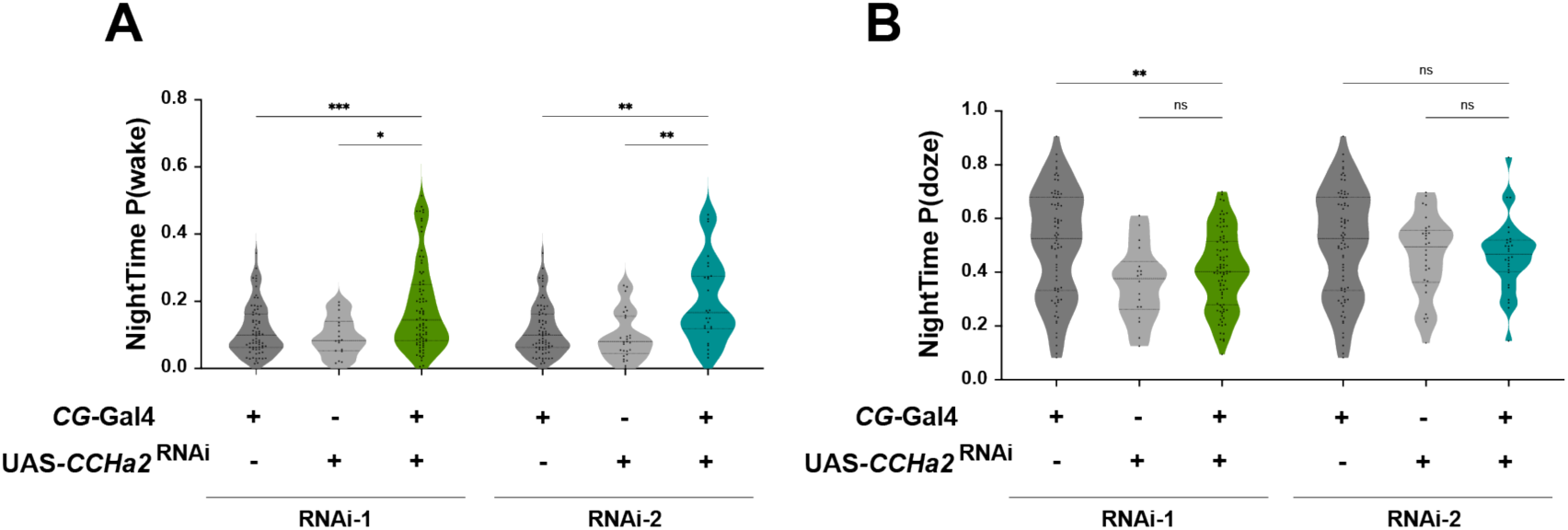
Nighttime sleep depth is altered in *CCHa2* knockdown flies. (A) The probability of waking (P(Wake)) during the night is significantly increased in *CCHa2* knockdown flies compared to controls (F_2,173_=7.622, *p* = 0.0007, F_2,128_= 6.035, *p* = 0.0031). (B) The probability of dozing (P(Doze)) during the night does not differ significantly across groups. (F_2,129_= 0.9135, *p* = 0.4037). Lines and error bars represent mean ± SEM.

**FIGURE S3.**
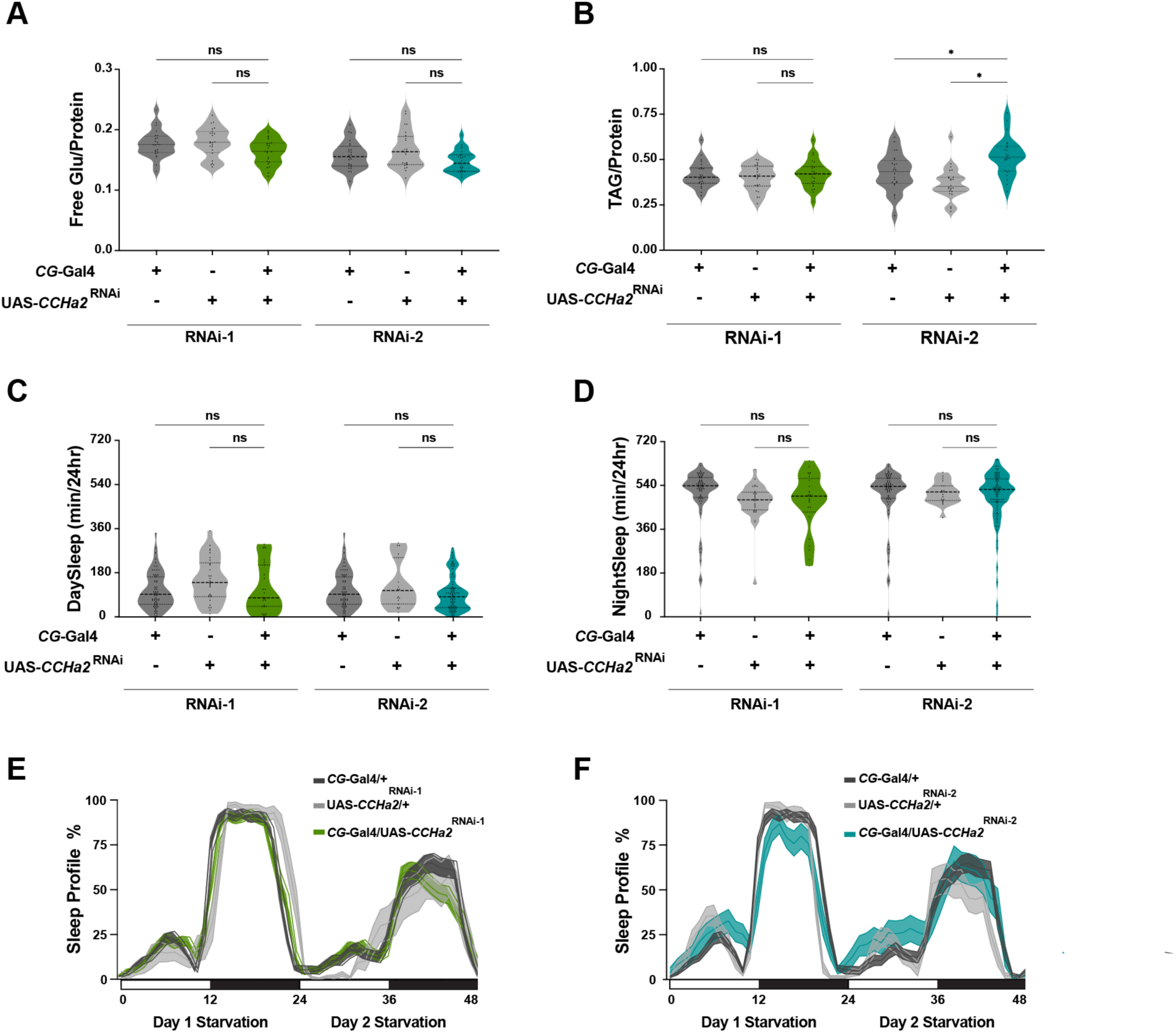
*CCHa2* knockdown impairs starvation-induced sleep suppression but does not affect energy storage. (A) Free glucose level, normalized to protein, show no significant differences in flies with *CCHa2* knockdown in the fat body in both RNAi lines (n=24, RNAi^1^: p= 0.7570, RNAi^2^: p= 0.0975). (B) Triglyceride level was only significantly different in the RNAi^2^ (n=24, RNAi^1^: p= 0.9571, RNAi^2^:p<0.01). (C-D) Sleep was recorded for 24 h under starvation (agar-only) conditions. Starvation suppressed sleep in control flies after starvation, but there was no change in *CG-*GAL4>*CCHa2*-RNAi flies as the result of starvation during daytime (N = 120, *p* = 2.941), and nighttime (N = 118, *p* = 0.2614). (E-F) Sleep profiles under starvation show no significant genotype differences (*p*= 0.9090)

**FIGURE S4.**
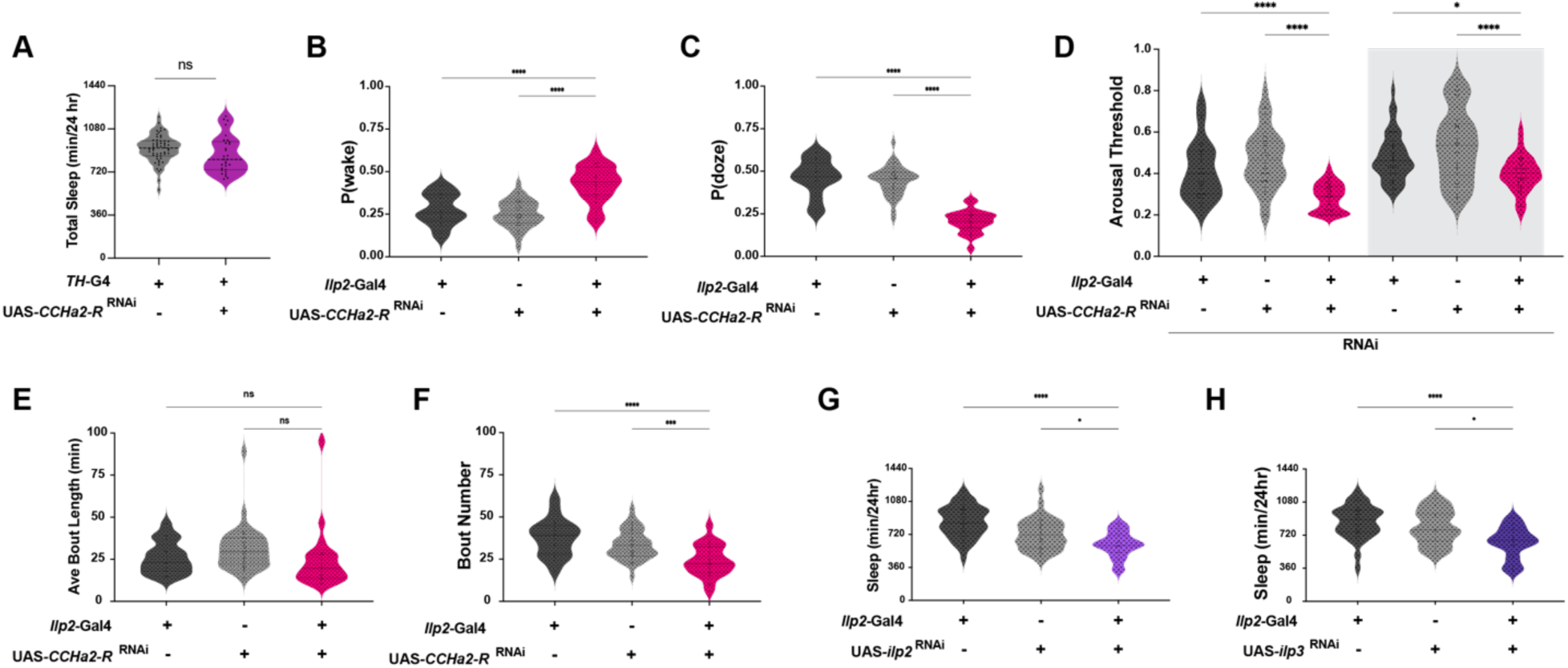
*CCHa2-R* and IPC-enriched genes modulate sleep and arousal in Drosophila. (A) *CCHa2R* knockdown under control of *TH-GAL4* has no effect on sleep (B-C) HMM analysis shows a significant increase in P(wake) (F _2, 75_ = 27.06, \*\**P* < 0.0001) and a significant decrease in P(doze) (F _2, 75_ = 63.63, *p* < 0.0001) in ilp2-Gal4>*CCHa2-R* knockdown flies. (D) Arousal threshold analysis reveals a marked reduction in sleep depth in CCHa2R knockdown flies compared to controls, F _5, 227_= 18.91, *p* <0.0001. (D-E) Targeted knockdown of *Ilp2* (F_2, 100_ = 13.47, *p* <0.0001) and *Ilp3* (F_2, 123_ = 20.84, \*\**p* <0.0001) using the IPC-specific driver *ilp2*-GAL4 significantly reduces total sleep compared to control. (G-H) Average bout length is unchanged, but the number of sleep bouts is significantly reduced in *CCHa2-R* knockdown flies.

