## Supplementary material for "The fat-body secreted neuropeptide CCHa2 signals insulin-producing cells in the brain to promote sleep": Table 1

| FBgn00 | BL/VDRC# | Gene | Total Sleep | Day Sleep | Night Sleep | ABL | ABN |
| --- | --- | --- | --- | --- | --- | --- | --- |
| FBgn0028645 | 55938 | beat-lb | 594.6666667 | 89.33333333 | 482.6666667 | 32.59798647 | 21.33333333 |
| FBgn0264001 | 43318 | bru3 | 795 | 331.4285714 | 462.1428571 | 30.84233123 | 29 |
| FBgn0265598 | 29454 | Bx | 582.6666667 | 271.6666667 | 309.6666667 | 16.31439019 | 37.53333333 |
| FBgn0264386 | 26251 | Ca-α1T | 612.8125 | 161.25 | 450.625 | 32.23969751 | 23.875 |
| FBgn0037963 | 28716 | Cad87A | 612.5 | 206.5625 | 404.6875 | 21.39089814 | 32.25 |
| FBgn0004580 | 28302 | Cbp53E | 660.3125 | 204.375 | 454.6875 | 26.96549039 | 28.875 |
| FBgn0033058 | 25855 | CCHa2-R | 532.8947368 | 111.5789474 | 419.7368421 | 22.30707293 | 30.18421053 |
| FBgn0033579 | 29419 | CG13229 | 757.5 | 286.4285714 | 470.3571429 | 31.11072137 | 33.42857143 |
| FBgn0050089 | 28321 | CG30089 | 735.3571429 | 258.2142857 | 475.7142857 | 24.97612434 | 32.5 |
| FBgn0052982 | 64555 | CG32982 | 604.0625 | 172.8125 | 429.6875 | 21.09832241 | 31 |
| FBgn0053543 | 64879 | CG33543 | 1005.3125 | 470 | 532.1875 | 56.36567621 | 20.375 |
| FBgn0085382 | 58291 | CG34353 | 718.125 | 237.8125 | 478.4375 | 32.49426141 | 24.0625 |
| FBgn0286778 | 36594 | CG46385 | 567.5 | 189.375 | 377.1875 | 15.79016185 | 36.5 |
| FBgn0031906 | 27660 | CG5160 | 677.3333333 | 281.6666667 | 395 | 22.61260866 | 34.73333333 |
| FBgn0262975 | 25984 | CnC | 736.5384615 | 218.0769231 | 517.6923077 | 47.73348489 | 25.38461538 |
| FBgn0005677 | 26758 | dac | 738.6666667 | 264.3333333 | 473.3333333 | 27.84665143 | 29.33333333 |
| FBgn0011274 | 30513 | Dif | 591.5625 | 133.4375 | 457.5 | 29.85637591 | 22.5625 |
| FBgn0265998 | 55908 | Doa | 665.625 | 225.3125 | 439.375 | 31.63983383 | 26.4375 |
| FBgn0005558 | 29339 | ey | 548.125 | 143.4375 | 404.375 | 22.78394479 | 23.375 |
| FBgn0000635 | 34084 | Fas2 | 704.375 | 290.9375 | 411.875 | 23.22061806 | 33.875 |
| FBgn0038197 | 32427 | foxo | 770.9375 | 310.3125 | 459.0625 | 24.05022267 | 34.25 |
| FBgn0004652 | 31593 | fru | 642.3333333 | 197 | 445.3333333 | 31.00943933 | 22.6 |
| FBgn0035245 | 67980 | GC | 694.6428571 | 218.2142857 | 475.7142857 | 31.22020161 | 24.57142857 |
| FBgn0036144 | 62221 | GIcAT-P | 877.75 | 361.75 | 512.5 | 24.5509305 | 38.95 |
| FBgn0001235 | 21655 | hth | 732.8571429 | 223.5714286 | 508.5714286 | 28.66414594 | 27.28571429 |
| FBgn0036046 | 32475 | Ilp2 | 625.2272727 | 221.3636364 | 399.0909091 | 19.05879935 | 35.72727273 |
| FBgn0044050 | 33681 | Ilp3 | 605.6603774 | 124.1891892 | 480.4054054 | 29.33520531 | 25.62264151 |
| FBgn0044048 | 33683 | Ilp5 | 763.6666667 | 274 | 488 | 27.48232388 | 29.4 |
| FBgn0285926 | 34977 | Imp | 623.5714286 | 173.9285714 | 448.9285714 | 26.55645071 | 26.07142857 |
| FBgn0026411 | 29341 | Lim1 | 714 | 298.333333 | 415.333333 | 23.7294316 | 19.8523417 |
| FBgn0265487 | 29585 | mbl | 748.6666667 | 255 | 493 | 23.22414123 | 37.26666667 |
| FBgn0261836 | 32377 | Msp300 | 584.6875 | 162.5 | 421.5625 | 21.95797473 | 27.625 |
| FBgn0002945 | 67788 | nkd | 622.8571429 | 156.4285714 | 466.4285714 | 28.099434 | 23.42857143 |
| FBgn0262614 | 28920 | pyd | 631.5625 | 182.5 | 448.4375 | 28.53021688 | 25.5 |
| FBgn0034157 | 53702 | resilin | 861.7647059 | 308.2352941 | 551.1764706 | 34.87695706 | 25.94117647 |
| FBgn0003255 | 31958 | rk | 725.9259259 | 247.7777778 | 476.6666667 | 26.53007068 | 32.18518519 |
| FBgn0261090 | 27295 | Sytβ | 842.8125 | 335 | 506.25 | 28.36682261 | 33.0625 |
| FBgn0086680 | 26228 | vvl | 677.8125 | 231.875 | 444.6875 | 26.15197715 | 29.125 |
| FBgn0036044 | 42874 | Zasp67 | 770.5 | 341.5 | 428 | 38.37842032 | 21.9 |
