## Supplementary material for "The fat-body secreted neuropeptide CCHa2 signals insulin-producing cells in the brain to promote sleep": Table 2

|  | BL/VDRC# |  | Total Sleep | Day Sleep | Night Sleep | ABL | ABN |
| --- | --- | --- | --- | --- | --- | --- | --- |
| FBgn0016122 | 67205 | Acer | 887.8125 | 234.6875 | 651.875 | 35.806038 | 28 |
| FBgn0267978 | 41673 | ap | 728 | 184 | 543 | 27.7076047 | 29.06666667 |
| FBgn0036449 | 25926 | Bmm | 595.625 | 92.5 | 503.125 | 33.0705881 | 20.25 |
| FBgn0034331 | 61348 | BombC2 | 1001.759259 | 277.2727273 | 563.8636364 | 33.1868177 | 27.90909091 |
| FBgn0038930 | 72072 | BomT3 | 712.8571429 | 168.5714286 | 541.4285714 | 35.0427239 | 23.10714286 |
| FBgn0038199 | 57562 | CCHa1 | 862.96875 | 251.5625 | 610 | 46.5509335 | 20.90625 |
| FBgn0038147 | v12830 | CCHa2 GD | 675 | 176.25 | 498.0357143 | 25.7268104 | 29.46428571 |
| FBgn0038147 | v102257 | CCHa2 KK | 666.744186 | 155.0641026 | 480.7692308 | 28.2309057 | 27.8974359 |
| FBgn0000276 | 64855 | CecA1 | 1082.89474 | 407.6315789 | 673.9473684 | 45.2743984 | 25.78947368 |
| FBgn0032836 | 64630 | CG10680 | 693.913043 | 175.8695652 | 515.6521739 | 32.2835386 | 25.56521739 |
| FBgn0039629 | 55617 | CG11842 | 1000 | 382.1153846 | 616.3461538 | 36.7944303 | 27.96153846 |
| FBgn0035496 | 53367 | CG14990 | 1077.38 | 214.1666667 | 551.1666667 | 34.8202645 | 24.66666667 |
| FBgn0030013 | 31118 | CG1583 | 756.304348 | 185.6521739 | 570.2173913 | 31.9711766 | 26.39130435 |
| FBgn0029990 | 65928 | CG2233 | 973.3333333 | 349.2592593 | 617.2222222 | 52.6131485 | 22.22222222 |
| FBgn0050280 | 53271 | CG30280 | 790.15625 | 204.53125 | 581.40625 | 37.455812 | 23.65625 |
| FBgn0031538 | 67017 | CG3246 | 670.2083333 | 169.1666667 | 496.4583333 | 28.2185523 | 26.66666667 |
| FBgn0052512 | 57437 | CG32512 | 1075.833333 | 449.5833333 | 624.5833333 | 50.4929993 | 25.875 |
| FBgn0053120 | 42934 | CG33120 | 882.8846154 | 281.9230769 | 627.1153846 | 47.0689623 | 21.38461538 |
| FBgn0035091 | 55245 | CG3829 | 764.677419 | 152.0967742 | 608.0645161 | 56.2342633 | 15.61290323 |
| FBgn0036622 | v109856 | CG4753 | 761.5625 | 128.4375 | 632.5 | 62.3490323 | 17.0625 |
| FBgn0039527 | 57376 | CG5639 | 1127.5 | 369.7727273 | 660.4545455 | 44.125658 | 25.45454545 |
| FBgn0038849 | 61174 | CG7079 | 1032.04545 | 222.0833333 | 597.0833333 | 45.5013246 | 19.875 |
| FBgn0036157 | 57161 | CG7560 | 824.1666667 | 222.0833333 | 597.0833333 | 45.5013246 | 19.875 |
| FBgn0027601 | 55272 | CG9009 | 711.25 | 115.625 | 595 | 48.3916576 | 18.4375 |
| FBgn0039805 | 57845 | cpr100A | 1052.894737 | 447.1052632 | 603.6842105 | 44.5524379 | 26.05263158 |
| FBgn0259938 | 29318 | Cwo | 1017.5 | 389.5833333 | 626.25 | 41.3828623 | 27.20833333 |
| FBgn0038095 | 67805 | Cyp304a1 | 809.6875 | 218.4375 | 589.375 | 32.2571027 | 28.75 |
| FBgn0000406 | 67351 | Cyt-b5-r | 1065.55556 | 417.2222222 | 646.1111111 | 76.0526608 | 16.22222222 |
| FBgn0033543 | 57486 | DaaO | 906.25 | 284.53125 | 620.625 | 41.8926383 | 22.3125 |
| FBgn0031461 | 34974 | Daw | 860.294118 | 264.1176471 | 595 | 29.8046493 | 30.35294118 |
| FBgn0010385 | 29524 | Def | 903.214286 | 294.6428571 | 607.8571429 | 31.14838 | 30.21428571 |
| FBgn0086687 | 35591 | Desat1 | 840.3125 | 255.46875 | 581.25 | 46.077809 | 20.15625 |
| FBgn0052185 | 62309 | Edin | 1080.625 | 366.8181818 | 565.9090909 | 49.7550621 | 21.63636364 |
| FBgn0003731 | 25781 | Egfr | 866.3636364 | 389.5833333 | 626.25 | 41.3828623 | 27.20833333 |
| FBgn0033483 | 55276 | Eiger | 718.055556 | 282.9032258 | 617.0967742 | 65.8135019 | 22.09677419 |
| FBgn0029769 | 34558 | frma | 905.483871 | 282.9032258 | 617.0967742 | 65.8135019 | 22.09677419 |
| FBgn0034199 | v15512 | GBP1 | 598.0434783 | 101.9565217 | 494.7826087 | 26.9499715 | 24.60869565 |
| FBgn0036428 | 42498 | Gbs-70E | 836.25 | 228.28125 | 607.03125 | 31.4660055 | 28.59375 |
| FBgn0032287 | 51867 | Gcst | 793.59375 | 206.5625 | 585.9375 | 37.352924 | 24.09375 |
| FBgn0001208 | 29540 | henna | 904.259259 | 330.5555556 | 572.4074074 | 31.9678786 | 33.07407407 |
| FBgn0001186 | 35155 | Hex C | 692.8125 | 302 | 494 | 56.2252483 | 28.2 |
| FBgn0040211 | 55629 | hgo | 729.6875 | 128.0555556 | 530.8333333 | 27.1873147 | 26.5 |
| FBgn0039297 | 57193 | Jhbp7 | 859.8 | 273.6 | 585 | 31.6689043 | 29 |
| FBgn0036992 | 52923 | Hpd | 659.722222 | 206.3461538 | 524.4230769 | 24.3294474 | 33.34615385 |
| FBgn0051092 | 54461 | LPR2 | 732.5 | 150.8695652 | 615.6521739 | 38.3589125 | 22.47826087 |
| FBgn0002563 | 27042 | LSP1 beta | 864.791667 | 292.5 | 571.875 | 28.2415225 | 34.04166667 |
| FBgn0002562 | 56039 | Lsp1-alpha | 760.47619 | 213.5714286 | 578.0952381 | 58.3772308 | 18.85714286 |
| FBgn0002565 | 57505 | Lsp2 | 872.142857 | 278.9285714 | 592.1428571 | 33.1675839 | 27.57142857 |
| FBgn0264691 | 34361 | Lst8 | 903.75 | 287.65625 | 611.40625 | 47.4436978 | 20.75 |
| FBgn0050359 | 41851 | MalA5 | 858.108108 | 344.5 | 537 | 31.9125037 | 32.15 |
| FBgn0050360 | 60398 | MalA6 | 806.25 | 293.9130435 | 547.3913043 | 29.5229122 | 33.17391304 |
| FBgn0033296 | 62252 | MalA7 | 528.4375 | 124.6875 | 403.75 | 17.4979261 | 33.09375 |
| FBgn0033297 | 55193 | MalA8 | 864.375 | 323.9583333 | 540.4166667 | 41.1874321 | 27.08333333 |
| FBgn0032381 | 55346 | MalB1 | 995.740741 | 405.7407407 | 590 | 36.13338 | 30.18518519 |
| FBgn0032382 | 62253 | malB2 | 904.62963 | 311.4814815 | 591.8518519 | 40.0239444 | 25.48148148 |
| FBst0041583 | 41563 | mt:srr | 888.333333 | 232.3809524 | 655.4761905 | 41.2381953 | 23.0952381 |
| FBgn0287423 | 29430 | NFP | 783.75 | 229.375 | 553.75 | 26.5807427 | 31.75 |
| FBgn0039801 | 67803 | NPC2h | 822.857143 | 140.7142857 | 682.1428571 | 49.056342 | 17.85714286 |
| FBgn0287423 | 29430 | Nplp2 | 745 | 155 | 500 | 18.8571429 | 35 |
| FBgn0013307 | 56103 | Odc1 | 801.25 | 197.34375 | 602.8125 | 46.4310803 | 19.78125 |
| FBgn0013308 | 60131 | Odc2 | 863.7096774 | 252.0967742 | 610.9677419 | 43.3855136 | 21.51612903 |
| FBgn0037146 | 51911 | p5cs | 723.75 | 333.6666667 | 622.6666667 | 57.471341 | 20.26666667 |
| FBgn0020513 | 62241 | Paics | 963.3333333 | 333.6666667 | 622.6666667 | 57.471341 | 20.26666667 |
| FBgn0036007 | 64029 | Path | 1076.07143 | 433.9285714 | 640.7142857 | 45.3892036 | 26.28571429 |
| FBgn0017558 | 28635 | pdk | 759.166667 | 193.3333333 | 564.8333333 | 28.4047132 | 28.3 |
| FBgn0003076 | v105820 | PGM1 | 831.37931 | 287.0689655 | 544.3103448 | 31.2541834 | 29.37931034 |
| FBgn0034909 | 61237 | Pippin | 1002.692308 | 390.8333333 | 607.5 | 48.5354664 | 22.41666667 |
| FBgn0041194 | 51492 | Prat2 | 819.53125 | 278.8888889 | 607.2222222 | 42.9955303 | 23 |
| FBgn0039593 | 62346 | Sid | 891.9444444 | 237.2222222 | 588.1481481 | 35.1964791 | 25.81481481 |
| FBgn0014031 | 51935 | Spat | 826.666667 | 237.2222222 | 588.1481481 | 35.1964791 | 25.81481481 |
| FBgn0033782 | 27026 | sug | 933.125 | 283.75 | 648.9583333 | 35.2029825 | 28.16666667 |
| FBgn0041180 | 67218 | Tep4 | 889.791667 | 277.7083333 | 610.4166667 | 32.2267538 | 30.45833333 |
| FBgn0023479 | 41890 | Teq | 732.5806452 | 160.8064516 | 571.2903226 | 37.9977558 | 21.48387097 |
| FBgn0261575 | 53379 | Tobi | 573.5714286 | 344.7619048 | 606.4285714 | 36.1291347 | 29.80952381 |
| FBgn0038838 | 65103 | TotB | 1052.659574 | 123.9583333 | 589.7916667 | 38.3592135 | 20.66666667 |
| FBgn0044812 | 18460 | TotC | 718.125 | 123.9583333 | 589.7916667 | 38.3592135 | 20.66666667 |
| FBgn0037607 | 32884 | Transketolase | 642.5 | 107.8125 | 534.6875 | 36.3589418 | 21.75 |
| FBgn0028978 | 41903 | trbl | 750.333333 | 190.3125 | 598.125 | 37.1631704 | 22.71875 |
| FBgn0003944 | 34993 | Ubx | 793.4375 | 190.3125 | 598.125 | 37.1631704 | 22.71875 |
| FBgn0030904 | 33988 | Upd2 | 732.8125 | 154.0625 | 582.1875 | 29.6574519 | 28.03125 |
